# Using the SSVEP to measure the SNARC-spatial attention effects in a parity judgment task

**DOI:** 10.1101/2023.04.11.536438

**Authors:** A. Mora-Cortes, R. Gulbinaite, K.R. Ridderinkhof, M X. Cohen

**Affiliations:** Life Science Department, Erasmus University College, Nieuwemarkt 1A, 3011HP, Rotterdam, The Netherlands; Laboratorio de Neurociencia y Comportamiento, Universidad de los Andes, Bogotá, Colombia; Lyon Neuroscience Research Center (CRNL), Brain Dynamics and Cognition Team, INSERM U1028, CNRS UMR5292, Université Claude Bernard Lyon 1, UdL, Lyon, France; Department of Psychology, University of Amsterdam, Weesperplein 4, 1018 XA Amsterdam, The Netherlands; Amsterdam Brain & Cognition (ABC), University of Amsterdam, Nieuwe Achtergracht 10, 1018 WB Amsterdam, The Netherlands; Radboud University and Radboud University Medical Center, Donders Institute for Neuroscience, Netherlands

**Keywords:** SNARC, spatial attention, parity judgment, SSVEP

## Abstract

Mental representation of numbers in the brain has been described as spatially organized on what is known as the “mental number line,” with small numbers to the left and large numbers to the right. This representation leads to the “SNARC effect” (Spatial-Numerical Association of Response Codes), which refers to (1) improved behavioral performance for “congruent” conditions (e.g., left-handed response to small numbers) as compared to “incongruent” conditions (e.g., right-handed response to small numbers) and (2) perceiving numbers induces an automatic shift in covert spatial attention. Nonetheless, there is scarce physiological evidence for (or against) the prediction that number magnitude induces an automatic shift of spatial attention. Here we recorded EEG during the bimanual SNARC-parity judgment task (classifying numbers as odd or even) in attempt to find electrophysiological evidence for number-magnitude-induced automatic shifts of spatial attention. Attention was measured using the SSVEP (steady-state visual evoked potential) from four task-irrelevant flickering frequencies (two to the left: 15 and 20 Hz and two to the right: 24 and 17.14 Hz) in combination with an optimal spatial filtering method. EEG analysis was performed on the flicker frequencies closest to the fixation point: 20 and 24 Hz for the left and right visual hemifields, respectively. We replicated the expected behavioral patterns predicted by the SNARC effect, such that participants performed better for congruent conditions than on incongruent conditions. We observed significant changes in the SSVEP amplitude with respect to the baseline for the left (20 Hz) and right (24 Hz) flicker for both the congruent and incongruent conditions. Statistically significant differences between the congruent and incongruent conditions were larger for the congruent conditions for the flicker stimuli on the left. Together, these findings support the hypothesis that when numbers are part of the task, but their magnitude is not relevant, the SNARC-spatial attention effect is elicited as a cognitive effect resulting from the mental representation of numbers and its relationship with the space representation, more than merely a motor effect.

## INTRODUCTION

Modern human innovations and advanced thinking like math and science have numbers as their cornerstone. Therefore, understanding how numbers are represented in the brain is a crucial topic for understanding human cognition and intelligence. An intriguing insight into the neural representation of numbers comes from the observation that numbers seem to be mentally represented on a so-called mental number line (MNL), where numbers are organized according to their increasing magnitude (Moyer & Landauer, 1967; Restle, 1970) (See Figure 1.1).

**Figure 1.**
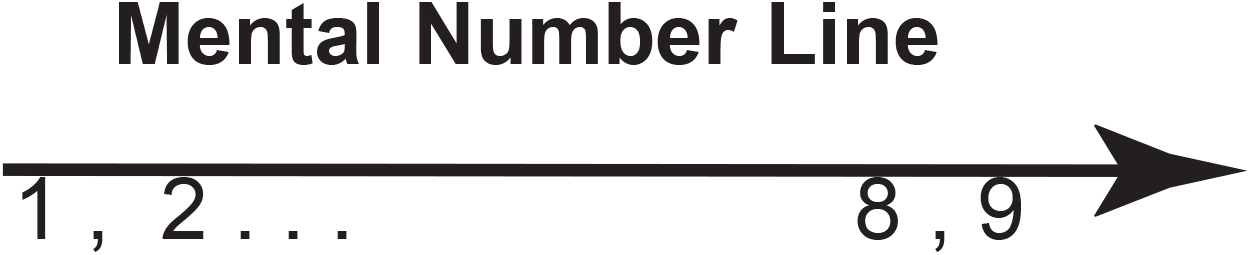
Sketch of number representation according to the MNL where small numbers (e.g., 1 and 2) are spatially represented to the left side and large numbers (e.g., 8 and 9) on the right side of the MNL.

Following this idea, Dehaene, Bossini & Giraux (1993) showed for the first time that numbers seem to be spatially represented in a horizontal MNL with a specific direction going from left-to-right according to number sign and magnitude. This means that small magnitude numbers like “1” or “2” are associated with the left side of space while large magnitude digits like “8” or “9” are associated with the right side of the space. The authors used the parity judgment task (classify odd vs. even numbers) across different experiments using different response mappings (e.g., even-left vs. odd-right, bimanual parity judgment) and concluded that participants were faster to respond to small-magnitude numbers, and slower to respond to large-magnitude numbers, with the left hand, and vice-versa for the right hand. The authors termed this the Spatial-Numerical Association of Response Codes (SNARC) effect. The SNARC effect is present not only when the number magnitude is relevant for the task (magnitude-judgment) but also when the magnitude information is not relevant (parity judgment; Fias & Fischer, 2004; Fias, Lauwereyns, & Lammertyn, 2001).

Although the SNARC effect was described in a bimanual response task, it has also been reported in pointing tasks (Fischer, 2003), eye-movements (Fischer, Warlop, Hill, & Fias, 2004; Schwarz & Keus, 2004), grasping (Andres, Ostry, Nicol, & Paus, 2008), verbal (Keus & Schwarz, 2005), and bipedal responses (Schwarz & Müller, 2006). Similarly, it has been shown that the SNARC effect is present in non-numerical tasks such as phoneme detection for digits’ names (Fias, Brysbaert, Geypens, & d’Ydewalle, 1996), and midpoint localization on long strings of numbers (Fischer, 2001). These studies show that the SNARC effect is a fundamental representation and is not linked to one particular response modality or experiment paradigm.

Supporting the notion that SNARC reflects the mental representation of numbers magnitude in this left-to-right spatial organization, it has been reported that perceiving numbers induces an *automatic shift of spatial attention* to the left when small numbers are observed and towards the right side of space when large numbers are perceived (Fischer et al., 2003). However, there have been difficulties in replicating the modulation of spatial attention induced solely by the mere observation of numbers, as described by Fischer, making its reliability difficult to ascertain.

On the one hand, several follow-up studies have replicated the SNARC effect (i.e., in parity or magnitude judgment tasks) and authors have claimed that it is explained by the MNL theory (Chinello, de Hevia, Geraci, & Girelli, 2012; Göbel, Calabria, Farnè, & Rossetti, 2006; Hesse & Bremmer, 2017; Müller & Schwarz, 2007; Schwarz & Keus, 2004; Shaki, Fischer, & Göbel, 2012; Weis, Estner, van Leeuwen, & Lachmann, 2016; Zohar-shai et al., 2017; Zorzi, Priftis, & Umiltà, 2002; See Fias & Fischer, 2005 for an extended review), while some others have proposed that the SNARC effect could be better explained by the stimulus response compatibility (SRC; Fitts & Seeger, 1953; Kornblum, Hasbroucq, & Osman, 1990), the dual-route cognitive model (Gevers et al., 2005; Gevers, Ratinckx, et al., 2006) or the polarity correspondence theory (PCT) of compatibility (Proctor & Cho, 2006). Results from fMRI (Pinel et al., 2004; Vogel, Grabner, Schneider, Siegler, & Ansari, 2013) and ERP studies (Gut et al., 2012; Keus, Jenks, & Schwarz, 2005) have supported this response target-stimulus related notion of the SNARC effect.

On the other hand, the spatial SNARC-spatial attention effect (to differentiate it from the classical SNARC effect observed in motor responses during parity or magnitude task) as described by Fischer et al. (2003) has been elusive to reproduce. While some authors have replicated the SNARC-spatial attention (Galfano et al., 2006; Ristic et al., 2006) claiming that this is not due to an automatic effect induced by numbers, other studies have failed to reproduce this SNARC-spatial attention effect (van Dijck, Abrahamse, Acar, Ketels, & Fias, 2014; Zanolie & Pecher, 2014).

This variability in the reported results suggest that mental representation of numbers, and that to elucidate the effect of numbers on allocation of spatial attention requires a direct measurement of attention while performing a task involving numbers. And despite the extended literature about the SNARC effect and the possible relation with allocation of spatial attention, most of the ERP or fMRI studies have focused on describing how numbers and mental representation of space interact to improve behavioral performance.

Similarly, some other studies have focused on explaining the cognitive components that would support such interaction between numbers and space inducing the SNARC effect. Nevertheless, in the SNARC literature there is a lack of evidence regarding the direct measure of spatial attention while performing a SNARC task. Therefore, in order to better understand whether numbers have a direct effect in allocating spatial attention either when it is explicitly or implicitly manipulated, direct brain measurements (as opposed to indirect behavioral measurements) may provide more insight if numbers are processed by our brain as cues for stimulus-driven spatial attention. Thus, to provide physiological evidence of the effect of numbers on spatial attention, and to test if the implicit manipulation of spatial attention by observing numbers would induce the SNARC spatial attention effect, in the present study we have combined a parity judgment task with direct brain measurement of spatial attention.

In this study we combined the steady-state visual evoked potential (SSVEP) technique with a parity judgment task as an alternative approach to provide further evidence that numbers induce spatial attention allocation while performing a SNARC task. The SSVEP has been extensively used to study attention because the amplitude of the SSVEP correlates with the amount of attention required by the task (Andersen & Müller, 2010; Müller, Teder-Sälejärvi, & Hillyard, 1998). The SSVEP is a rhythmic neural response that tracks a flickering sensory stimulus (a.k.a. photic drive; Luck, 2005; Regan, 1989) and can be induced with a flickering stimulation in the range from 1 to 100 Hz (Herrmann, 2001).

On the other hand, the parity judgment task allows evaluating the effects of allocation of spatial attention associated with the implicit expectation of numbers in the task (Doherty et al., 2005; Posner, 1980; Posner et al., 1980).

Thus, the main goal of this study was to use the SSVEP method to measure whether numbers elicit an *automatic shift of spatial attention* during a SNARC task. To do so, we combined the parity task with the SSVEP where the numbers, but not their magnitude, were relevant for the task: implicit attention. Because we did not explicitly manipulate spatial attention in our task, we could evaluate whether the simple observation of numbers would reveal an attention-modulation effect for four different flicker frequencies (15 Hz, 20 Hz, 24 Hz and 17.14 Hz). We hypothesized that numbers would induce the SNARC-spatial attention effect even though their magnitude is not relevant for the task. Therefore, we predicted that EEG data would reveal a SNARC-spatial attention modulation effect on the SSVEP amplitude for the different flicker frequencies, with larger effects for conditions with small-odd or large-even numbers for the flicker frequencies (because these are the furthest apart congruent conditions from the set of numbers used in our task) placed to the left and right of the visual field, respectively.

## METHODS

### Participants

Thirty subjects participated in the experiment. During EEG pre-processing, the data from one subject was rejected due to excessive eye-blink and artifacts in more than 30% of the trials. The data from the other 29 participants (12 males, mean age 25.38, two left-handed) were included in the final analysis. All subjects had normal or corrected-to normal vision. The study was conducted in accordance with the Declaration of Helsinki, relevant laws, and institutional guidelines, and was approved by the local ethics committee at the Psychology Department at the University of Amsterdam.

Participants signed an informed consent document prior to the beginning of the experiment, and they were paid for their participation. Subjects are the same who participated in the target detection task, and they had a break of 15 min between experiments. The order of the experiments was counterbalanced between subjects.

### Task

We designed our task to evaluate the attention-modulation of the SNARC effect on the SSVEP amplitude in a parity judgment task for congruent (small numbers: left button and large numbers: right button) and incongruent (small numbers: right button and large numbers: left button) conditions. Participants were seated at a distance of approximate 90 cm from and at the level eye of a 24’’ BenQ monitor (1,920 × 1,080 pixels) with a refresh rate of 120 Hz. Stimulus presentation and behavioral response collection were controlled by custom-written scripts in Matlab (The Mathworks, Natick, MA) using the Psychophysics Toolbox (Brainard, 1997; Kleiner et al., 2007; Pelli, 1997).

Participants were requested to direct their gaze to the fixation point in the center of the screen (approximately 0.39° of visual angle -v.a.-; note that all measurements are approximate because subjects were not restrained e.g., by a chin rest) during the entire task. Participants were informed that numbers (0.86° v.a.) would be displayed on top of the fixation point and that they should press the left-mouse button to odd numbers and the right-mouse button to even numbers. This response mapping was not counterbalanced across subjects or experimental blocks. Participants were requested to respond as quickly and accurately as possible after stimulus presentation and they did not receive feedback on their performance. Figure 2.2 shows the timeline of a trial sequence presented to each participant.

**Figure 2.**
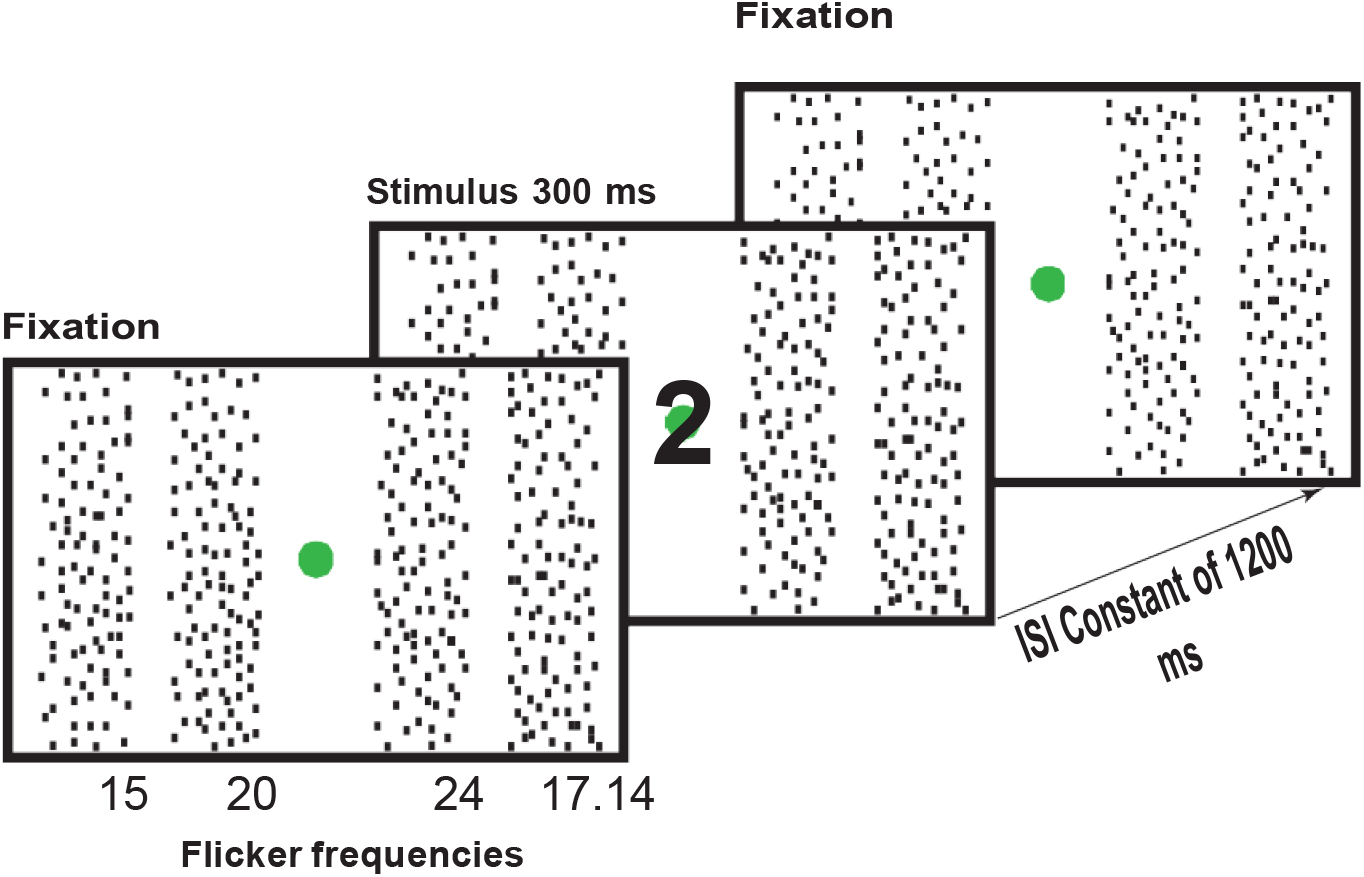
Example of the SNARC parity judgment task used in this study. Here an exampleof the individual timeline presentation of one trial. The flicker frequencies in the *x* axis (15, 20, 24 and 17.14 Hz) correspond to the flicker rates in Hz for the corresponding columns.The presentation and the flicker rate for each column were constant throughout each block (see text for details).

On each side of the fixation point (1° v.a. from fixation) two columns of flickering dots (0.11° v.a. wide each dot) were presented and participants were requested to ignore these flickering dots since they were not relevant for the task. The four columns flickered at a particular frequency: 15 Hz to the leftmost side of fixation and 20 Hz to the left side of the fixation point; and 24 Hz to the right of the fixation point and 17.14 Hz to the rightmost side of fixation. The columns were 1° v.a. apart from each other. And both the fixation point and the flickering columns were presented continuously throughout each block. Square-wave times series were created for each frequency (50% on per frequency).

In each block, numbers were presented for 300 ms followed by an interstimulus interval (ISI) of 1200 ms and the ISI was the same on all blocks. The stimuli numbers were 1,2,3,4,6,7,8 and 9 and they were separated in two groups in such a way that on each block two even and two odd numbers were presented, and the total amount of even and odd numbers displayed were equal on each block. Although the same number could be presented at most two consecutive times, each group of numbers was presented in pseudo-randomized order with the restriction that the same group of numbers was not presented consecutively. Then each participant completed nine blocks, each comprising 79 trials lasting approximately 2 minutes. Self-paced breaks were given between blocks and the full session lasted 20 minutes.

Finally, subjects were visually and verbally instructed and performed a practice block of 79 trials before starting the actual experimental session.

### Behavioral data analysis

Trials with RTs faster than 150 ms or slower than 2.5 standard deviations of the mean per condition and all error trials were excluded from behavioral analysis. From each participant, we computed number magnitude (small and large) and congruency (congruent for small odd numbers and large even numbers and incongruent for large odd numbers and small even numbers) obtaining four experimental conditions: small congruent (Scong), small incongruent (Sinc), large congruent (Lcong) and large incongruent (Linc). The statistical analyses of the four experimental conditions were performed using the statistical software SPSS 24.0 (SPSS, Chicago, IL).

The analyses for the differences between conditions were tested using repeated measures analyses of variance (ANOVA) with factors: number magnitude (for small: 1,2,3,4 and large: 6,7,8,9 numbers) and congruency (congruent and incongruent). Behavioral responses for accuracy were collected as correct or incorrect. Correct responses were defined as the response within the defined time window for RTs and the response button corresponding to the number displayed (left for odd and right for even numbers). Target presentation without response or responded with the incorrect button (left for even or right for odd numbers) were defined as incorrect response. However, because accuracy was defined as a binary response (right or wrong) only right responses were included for further analysis.

In cases of violation of sphericity, significance levels were adjusted using the Greenhouse-Geisser correction when performing repeated-measures ANOVA, and all the effect sizes are reported as partial eta-squared (η_p_^2^) or Cohen’s *d*, when appropriate.

### EEG recordings and preprocessing

The EEG data were recorded with a sampling rate of 1024 Hz from 64 scalp electrodes. Two additional electrodes were placed in the outer canthi of each eye to measure horizontal eye movements (HEOG). All data were collected using a BioSem Active Two system (see www.biosemi.com for details). Data were preprocessed with the EEGLAB toolbox for Matlab (Delorme & Makeig, 2004).

The data were high-pass filtered offline at 0.5 Hz and re-referenced off-line to the scalp average. After baseline subtraction (−200 to 0 ms pre-stimulus base line), trials with muscle and blinking artifacts during the stimulus were visually identified and manually removed (average of 6%). Then, additional artifacts or any kind of noise not related with brain activity were removed using independent component analysis in EEGLAB (Delorme & Makeig, 2004). All the EEG analyses involved only correct trials.

### SSVEP analysis

We performed two sets of analyses on the SSVEP data for each of the flickering frequencies used in the experiment: “static SSVEP”, in which the fast-Fourier transform (FFT) was computed over the full-time window of each trial (0 to 1500 ms), and “dynamic SSVEP”, in which a narrow band-pass Gaussian filter was applied to inspect the time-varying changes in SSVEP amplitude. Whereas the former method provides higher frequency resolution and increases the signal-to-noise (SNR) characteristics, the latter method provides higher temporal precision and thus the ability to detect time-varying changes in SSVEP amplitude resulting from phasic tasks events. The time windows for analysis were applied equally to all conditions.

SSVEP was extracted using a spatial filtering method termed rhythmic entrainment source separation (RESS; Cohen & Gulbinaite, 2017). The RESS method involves using generalized eigendecomposition to construct an optimal spatial filter that maximizes the SSVEP effect. The eigenvector with the largest eigenvalue maximally differentiates two covariance matrices (a “signal” and a “reference” covariance matrix), and that vector is used as channel weights to combine data from all electrodes. The signal covariance matrix was obtained from the narrowband-filtered signal at each stimulus frequency. The reference covariance matrix was obtained from data filtered at the closely spaced up- and down-neighbor frequencies. Importantly, the spatial filters were computed per frequency pooling data across all conditions; thus, although the spatial filter maximizes the overall SSVEP amplitude, there is no potential for biasing any condition comparison. The power spectrum was computed via the FFT for each flicker condition (15 Hz, 20 Hz, 24 Hz and 17.14 Hz), pooling over magnitude and congruency conditions over the entire trial time window. The static SSVEP was computed as the SNR as the ratio of the power at the flickered frequency to the average of neighboring frequencies surrounding each frequency (± 2 Hz, excluding a 0.5 Hz window around each center frequency) allowing us to convert power spectrum into SNR units to facilitate comparability across groups. See Figure 3 for an example of the topographical maps and SNR spectra obtained with RESS spatial filter.

**Figure 3.**
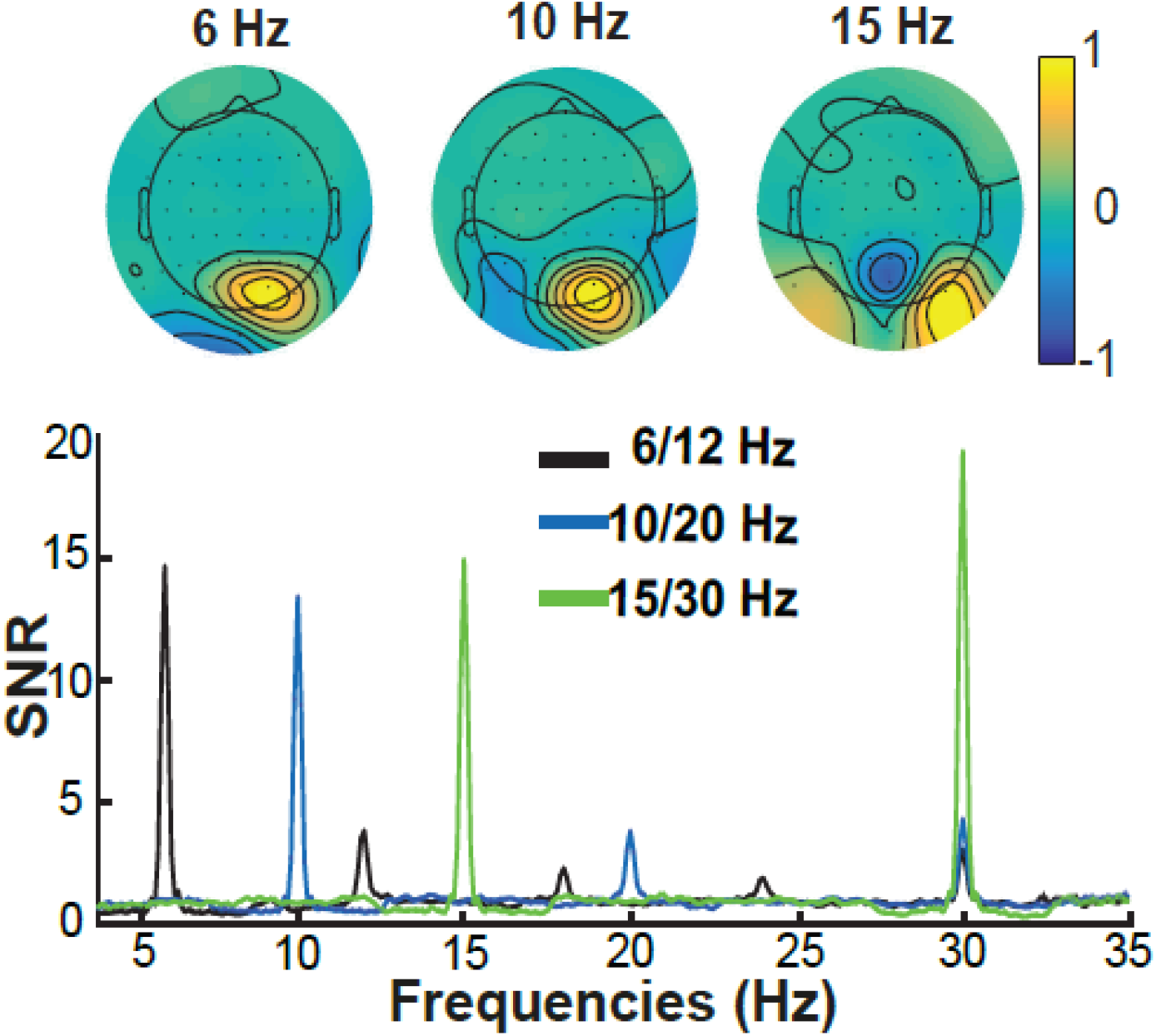
Topographical maps and power spectra from participant 17 as illustration. Top panel represents the RESS topographical maps for the frequencies used in our experiment showing higher SNR. Bottom panel shows the corresponding power spectrum at the corresponding fundamental frequency (6, 10, 15Hz) and the second harmonic (12, 20, 30 Hz).

For the computation of the dynamic SSVEP amplitudes, a narrow-band Gaussian filter was implemented for each of the flicker conditions with a full width at half-maximum (FWHM) of 6 Hz (the standard deviation of the filter; while a narrow filter provides better spectral precision the temporal precision is reduced, therefore we use a high FWHM to increase the temporal resolution of the filter). This Gaussian was point-wise multiplied by the power spectrum of the EEG data, and the inverse Fourier transform was applied to reconstruct the time course of the activity (i.e., convolution). After filtering the data, the amplitude of the frequencies of interest was extracted as the square magnitude of the result of the Hilbert transform. We call this time series the “SSVEP amplitude time series”. To facilitate comparisons across subjects, the SSVEP time series were baseline-normalized to a prestimulus period of -600 to -200 ms, separately for each frequency and condition specific. Note that in our task the flickering stimulus was present all the time, including the peri-stimulus period, thus the baseline was computed using the average of all conditions within each frequency, allowing for detection of possible condition differences at peri- and post-stimulus activity. Baseline normalization allows rescaling the activity relative to the selected baseline, which facilitates comparison between frequencies (Cohen, 2014).

### SSVEP statistical analysis

An initial visual inspection of the SSVEP results was performed for all the subjects and all the four flickering frequencies (15 Hz, 20 Hz, 24 Hz and 17.14 Hz) to confirm the SSVEP effects of the experimental paradigm. However, the statistical analyses were performed for the flickering stimuli closer to the fixation point (20 Hz and 24 Hz), first, because the analyses were focused on observing differences between the different experimental conditions and not between different flickering frequencies, and these two frequencies were optimal for further analysis; second, after performing initial analyses we didn’t observed significant effects for the 15 Hz and 17.14 Hz, and because these two frequencies were used as control conditions for the attention-modulation effects on the task. The SNR analyses for the differences between conditions were tested using repeated measures ANOVA with factors number magnitude (small and large), frequency (20 Hz and 24 Hz) and congruency (congruent and incongruent). In order to detect time-varying changes in SSVEP amplitude resulting from phasic task events, statistical analyses of the EEG SSVEP amplitudes time series were implemented for these eight experimental conditions (number magnitude, flicker and congruency). Nonparametric permutation testing was used in combination with cluster correction to evaluate condition differences in dynamic SSVEP over time (Maris & Oostenveld, 2007). To create a distribution of null-hypothesis time courses, the difference of the congruent – incongruent conditions for each frequency and number magnitude factor, for each subject was computed, and then multiplied this condition difference for a random subset of subjects by -1 (which is equivalent to randomizing incongruent – congruent vs. congruent – incongruent). This was repeated for 1,000 iterations. Finally, cluster correction was implemented to correct for multiple comparisons over the all-corresponding time points of the selected time window with a threshold value for pixel and cluster-levels of *p* = .05.

## RESULTS

### Behavioral results

Here we report only the relevant significant ANOVA results; see Table 1 for descriptive statistics and additional inferential statistics. For the RTs significant effects were observed for the number magnitude (*F*(1,28) = 21.05, *p* < .001, η_p_^2^ = .429), and congruency (*F*(1,28) = 18.80, *p* < .001, η_p_^2^ = .402) factors and the interaction between number magnitude and congruency (*F*(1,28) = 13.36, *p* = .001, η_p_^2^= .323) meaning that participants were faster in the congruent conditions (*M* = 465.55 ms, *SE* = 10.59) as compared to the incongruent conditions (*M* = 485.31 ms, *SE* = 11.31) (Figure 4.3). Similarly, participants observed faster responses in trials presenting numbers with small magnitude (*M* = 468.98 ms, *SE* = 10.40) as compared when the stimulus was a large magnitude number (*M* = 481.87 ms, *SE* = 11.20) (Figure 4.a).

**Table 1.** Behavioral statistical results from all the conditions

|  |  | Reaction times (ms) |  | Accuracy (%) |  |
| --- | --- | --- | --- | --- | --- |
| Number magnitude | Congruency condition | Mean | SD | Mean | SD |
| Descriptive statistics |  |  |  |  |  |
| Small | Congruent | 488.68 | 54.28 | 97.80 | 2.86 |
|  | Incongruent | 489.28 | 64.15 | 94.89 | 3.32 |
| Large | Congruent | 482.41 | 64.32 | 97.01 | 3.14 |
|  | Incongruent | 501.33 | 61.24 | 94.70 | 4.78 |
| Variable |  | df | F | p* |  |
| Repeated measures ANOVA for reaction times |  |  |  |  |  |
| Number magnitude |  | 1 | 21.050 | .000 |  |
| Congruency |  | 1 | 18.796 | .000 |  |
| Number magnitude X Congruency |  | 1 | 13.359 | .001 |  |
| Repeated measures ANOVA for accuracy |  |  |  |  |  |
| Number magnitude |  | 1 | 1.141 | .295 |  |
| Congruency |  | 1 | 12.417 | .001 |  |
| Number magnitude X Congruency |  | 1 | .346 | .561 |  |

Note: Upper table contains the descriptive statistics for RTs and accuracy. Middle and bottom tables contain the ANOVA results for RTs and accuracy, respectively. SS = sum of squares; MS = mean square. \* $p < .05$

**Figure 4.**
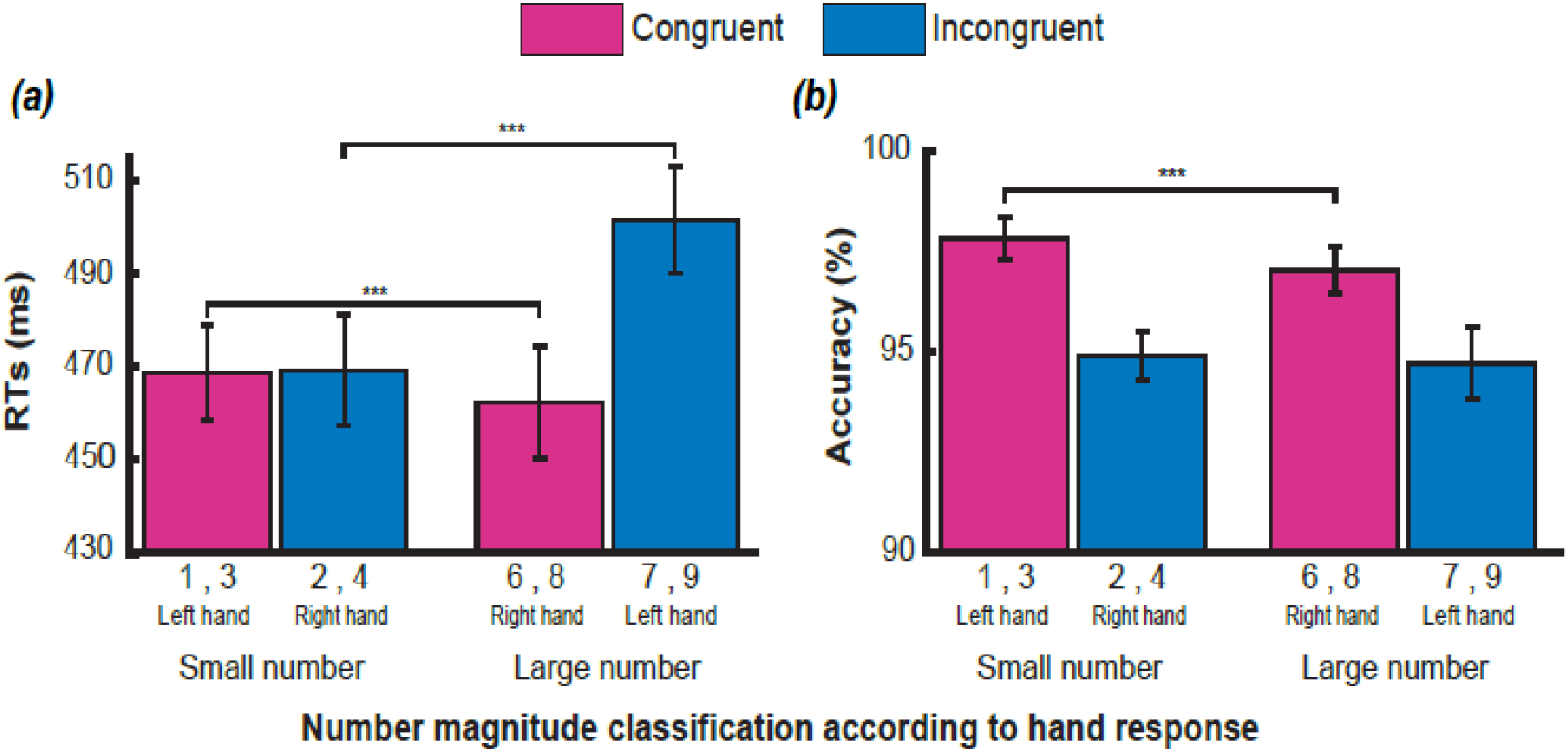
Behavioral responses parity judgment task (odd-left hand, even-right hand). a) RTs across subjects. Significant effects for the magnitude number (small/large) and congruency (congruent/incongruent) factors were observed. Participants showed faster response for congruent conditions as described by the SNARC effect. b) Accuracy was higher for congruent vs. incongruent trials. And better performance was observed for congruent trials with small numbers as compared with congruent trials presenting large numbers. Note that this improvement in performance was not statistically significant (number magnitude factor *p* > .1). Error bars represent standard error of the mean. ^***^ *p* < .001

For all the conditions, accuracy was above 90%. Congruent trials showed a significant effect as compared with incongruent trials (*M* = 97.4, *SE* = 0.5 and *M* = 94.8, *SE* = 0.6; *F*(1,28) = 12.42, *p* = .001, η_p_^2^ = .307) (Figure 4.3b), meaning that participants were more accurate classifying the stimulus number as odd or even when the response button was in the same side as the position of the number (e.g., small numbers with the left hand and large numbers with the right hand) as compared with trials where the response button was in the opposite side of the number and its mental representation (e.g., small numbers with the right hand and large numbers with the left hand). The results for the effects of number magnitude factor and the interaction between congruency and number magnitude were not significant. Because the response mapping was not counterbalanced across subjects or experimental blocks, we analyzed the individual performance regarding the hand response and number magnitude. With this we wanted to 1) confirm our group behavioral results and 2) provide additional behavioral evidence for the presence of the SNARC effect regarding the not counterbalance response mapping. Repeated-measures ANOVA for subjects average with factors hand (left or right) and number magnitude (small or large) showed significant effects for the hand (*F*(1,28) = 14.05, *p* = .001, η_p_^2^ = .335) and number magnitude (*F*(3,84) = 17.578, *p* <.000, η_p_^2^ = .386) factors as well as for the response hand X magnitude interaction (*F*(3,84) = 20.87, *p* < .000, η _p_^2^ = .427). These results support the SNARC effect we observed in our task (Figure 5). Altogether, our behavioral results showed that we could replicate the SNARC effect in the parity judgment task as has been described in previous studies, supporting the target-stimulus related notion of the SNARC effect.

**Figure 5.**
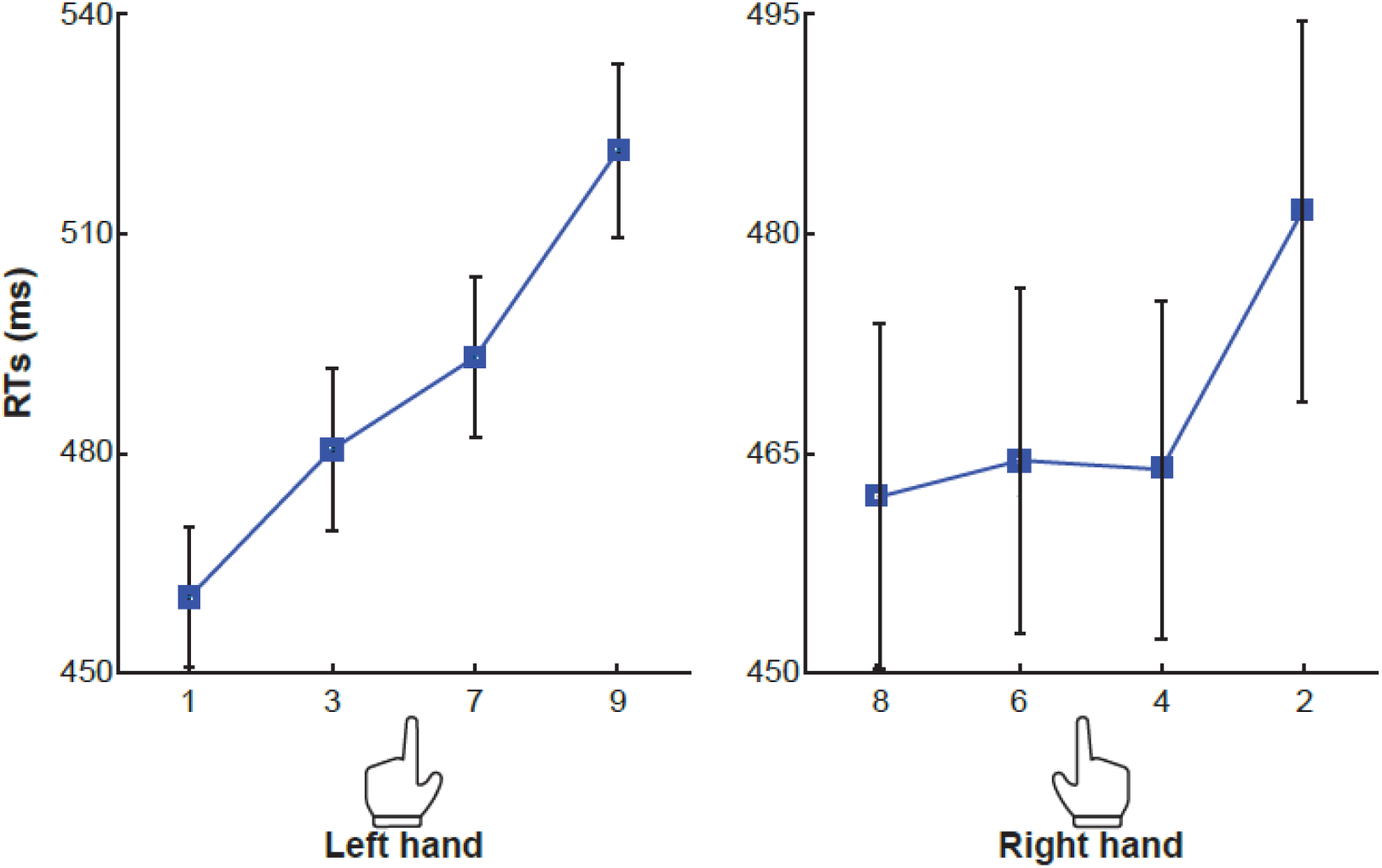
RTs obtained by number and response hand across subjects. The SNARC effect was observed for each number according to the button response: faster RTs with the left hand for 1 and 3 and for the right-hand numbers 8 and 6. Error bars represent standard error of the mean.

### EEG results

#### SNR results

We focused our EEG SSVEP analyses in the 20 Hz and 24 Hz flicker frequencies (see Methods). The SNR spectra exhibited robust peaks at both of these flicker frequencies, indicating that the design of our experiment was appropriate to elicit robust SSVEP effects during parity judgment task. The group average results showed RESS topographical maps with high SNR for the 20 Hz and 24 Hz. And high-power spectrum at the corresponding frequencies was also observed in the group average for both experimental conditions: congruent (small number/left button, large numbers/right button) and incongruent (small numbers/right button, large numbers/left button) (Figure 6).

**Figure 6.**
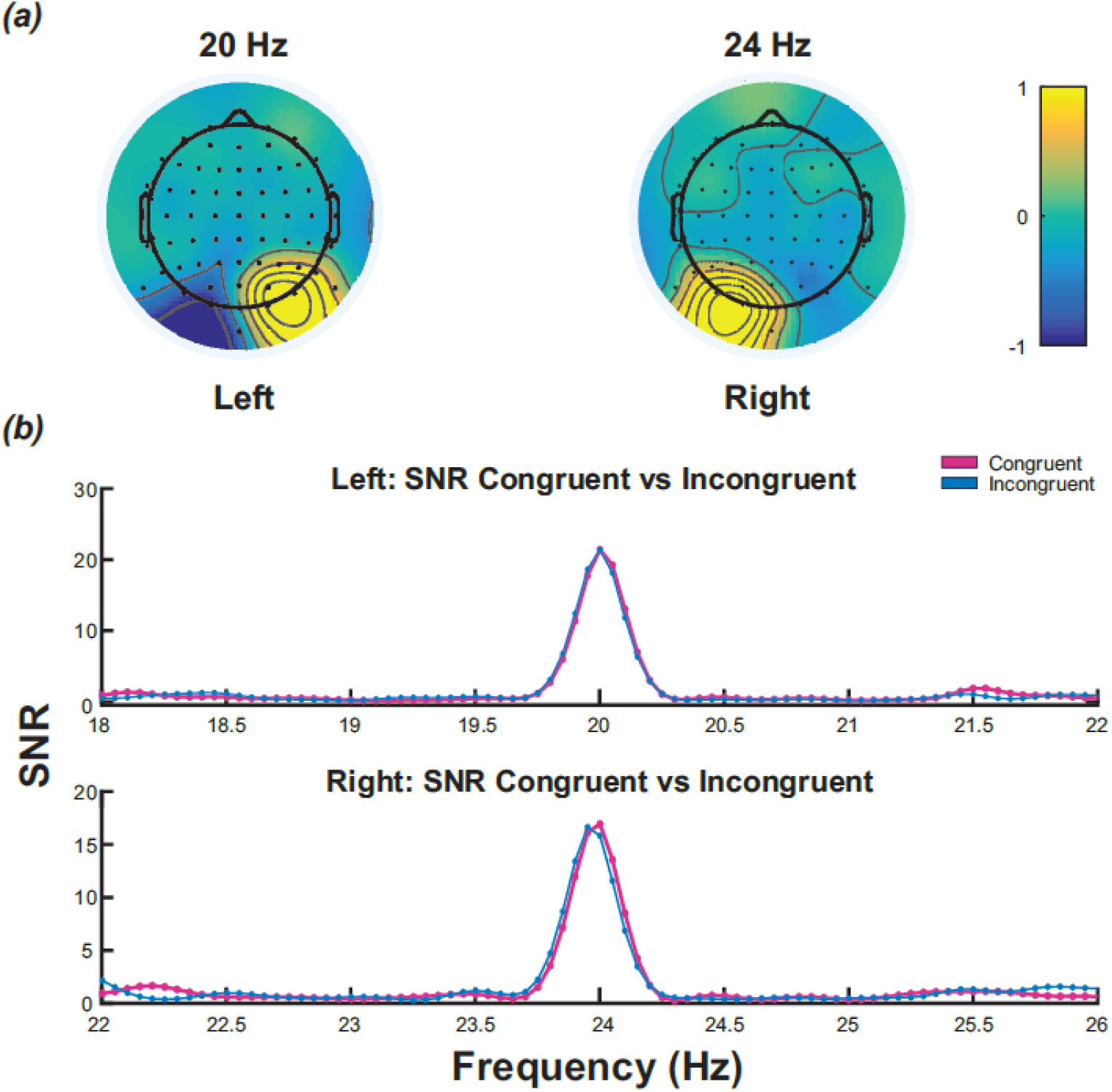
Group average SSVEP results. (a) Topographical maps across subjects for the flicker stimulus closest to the fixation point (20 Hz to the left, 24 Hz to the right). (b) Average SNRs per frequency (20 Hz top, 24 Hz bottom panel) for the congruent and incongruent conditions. Values in the *y* axis correspond to converting power spectrum to SNR units.

Our key hypothesis was that observation of numbers would produce the SNARC-spatial attention effect during the parity judgment task, therefore we expected SSVEP SNRs to be higher in 20 Hz (placed to the left of fixation) for small magnitude numbers and 24 Hz (placed to the right of fixation) for large magnitude numbers in the congruent conditions.

Repeated-measures ANOVA (with factors number magnitude, congruency and frequency) showed significant main effects for the frequency factor (*F*(1,28) = 18.87, *p* < .001, η _p_^2^ = .403). However, neither the main effect for congruency (*F*(1,28) = .002, *p* > .965, η _p_^2^ = .000), the number magnitude (*F*(1,28) = 2.98, *p* >.095, η _p_^2^ = .096) nor any of the interactions involving magnitude number, all *p’s >*.1 were significant. Figure 7 shows bar plots of the SNR for the 20 Hz and 24 Hz flicker frequencies for both congruent and incongruent conditions.

**Figure 7.**
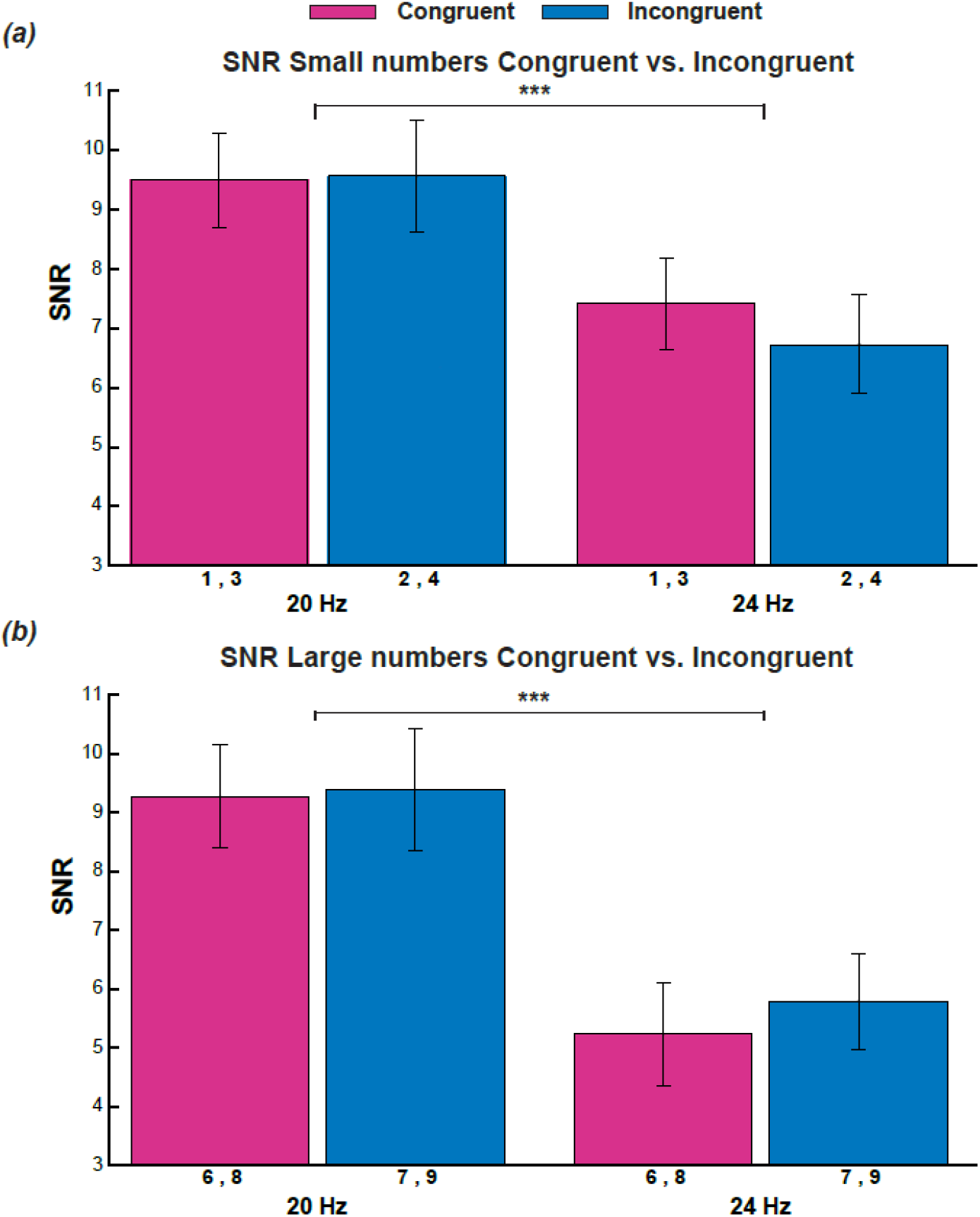
Statistical main effects were obtained for the frequency factor for the SSVEP SNR. Note that SNR values were higher for the flicker frequency placed to the left of the fixation point (20 Hz) as compared with the flicker stimuli to the right (24 Hz) for trials with (a) small or (b) large magnitude numbers (See figure 4.2). Values in the *y* axis correspond to converting power spectrum to SNR units. Error bars represent standard error of the mean. ^***^ *p* < .001

Higher SNRs can be observed for the flicker placed to the left of fixation (20 Hz) for both the congruent and incongruent condition with small (a) and large numbers (b) as compared with the flicker placed to the right of fixation (24 Hz). Also, in the upper panel it can be observed that for the flicker on the right the SNR was higher for the congruent conditions when displaying small numbers (i.e., numbers 1 and 3) as compared with incongruent trials (i.e., numbers 2 and 4), this large difference between the congruent and incongruent conditions was not observed for the flicker on the left. On the other hand, in the lower panel it can be observed that for the right flicker the SNR for congruent conditions with large numbers (i.e., 6 and 8) showed smaller SNR as compared with the incongruent conditions with large numbers (i.e., 7 and 9).

Although we did not obtain any statistically significant effects involving the magnitude factor, we performed a post hoc analysis in which we compared the SNRs between small vs. large numbers where we computed the average of the small numbers for the left (20 Hz) and right (24 Hz) flicker compared to the average of the large numbers. Paired samples *t* test showed no significant effects between small and large numbers (all *p’s* > .1). Table 4.2 contains all the ANOVA and *t* test results for the left (20 Hz) and right (24 Hz) flicker SNRs.

**Table 2.**
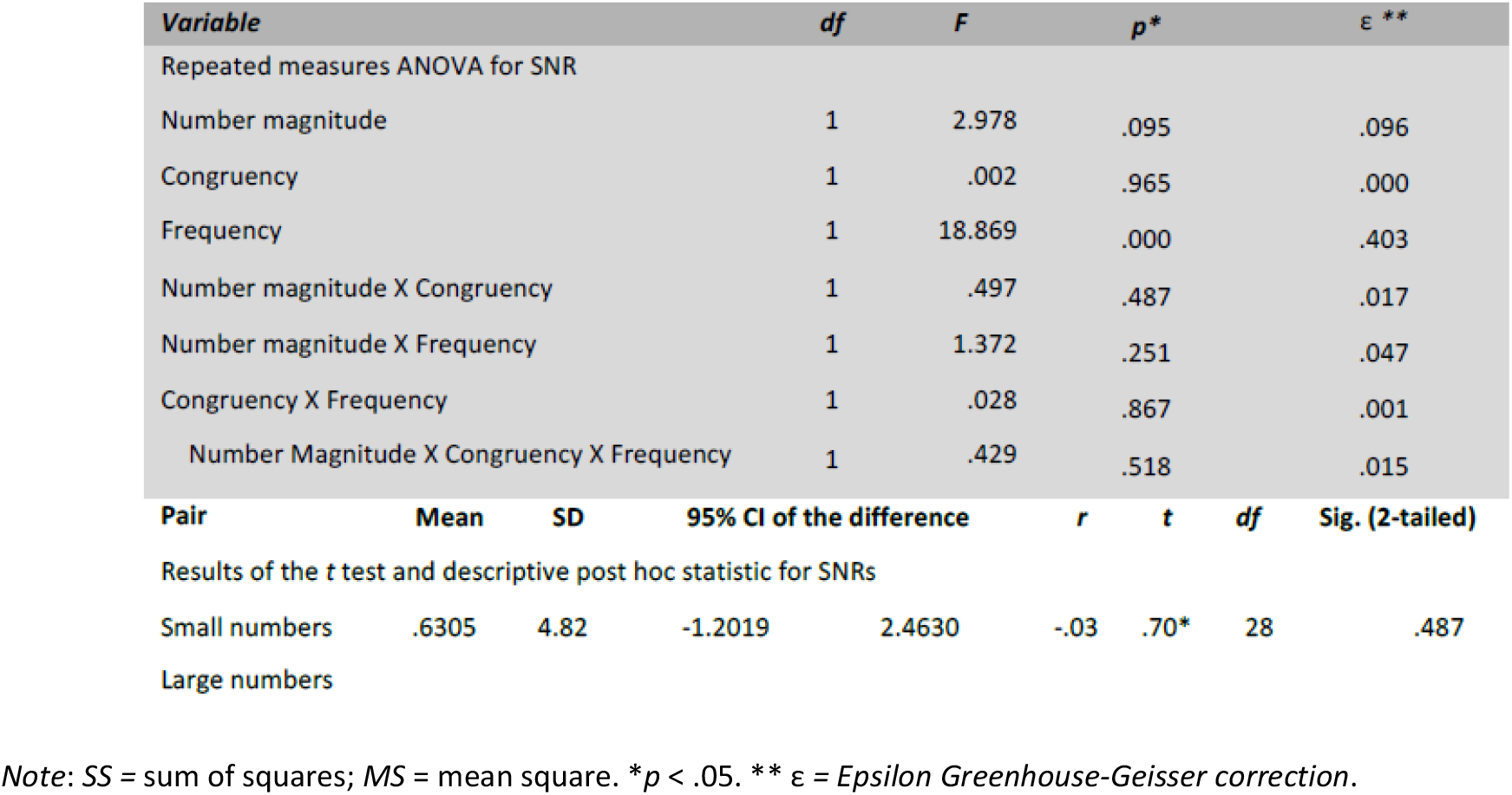
SNR Repeated measures ANOVA

#### Dynamic SSVEP results

The static SSVEP exhibited high SNR, but it is possible that the *automatic shift of spatial attention* was transient, and therefore it might not have been visible in the static analysis, which requires a long-time window for spectral resolution. Therefore, we applied a dynamic SSVEP amplitude analysis to search for temporally localized modulations. Our key hypothesis was that observation of numbers would produce the SNARC-spatial attention effect during the parity judgment task, therefore, we expected SSVEP amplitudes to be higher for the flicker on the left (20 Hz) and on the right (24 Hz) for the small and large magnitude numbers, respectively.

To facilitate comparison between frequencies we performed the percentage change baseline normalization to a pretrial baseline of -600 to -200 ms (See Methods). Because the flicker stimulation was continuous over time, meaning that any change relative to baseline is relevant, we first evaluated whether each flicker condition deviated from 100% (i.e., baseline). Therefore, we performed a t-test for the difference between every time point from each of the flicker condition against 100 (all *p* values < .05, cluster-corrected for multiple correlated tests over time points, see Methods and Figure 8).

**Figure 8.**
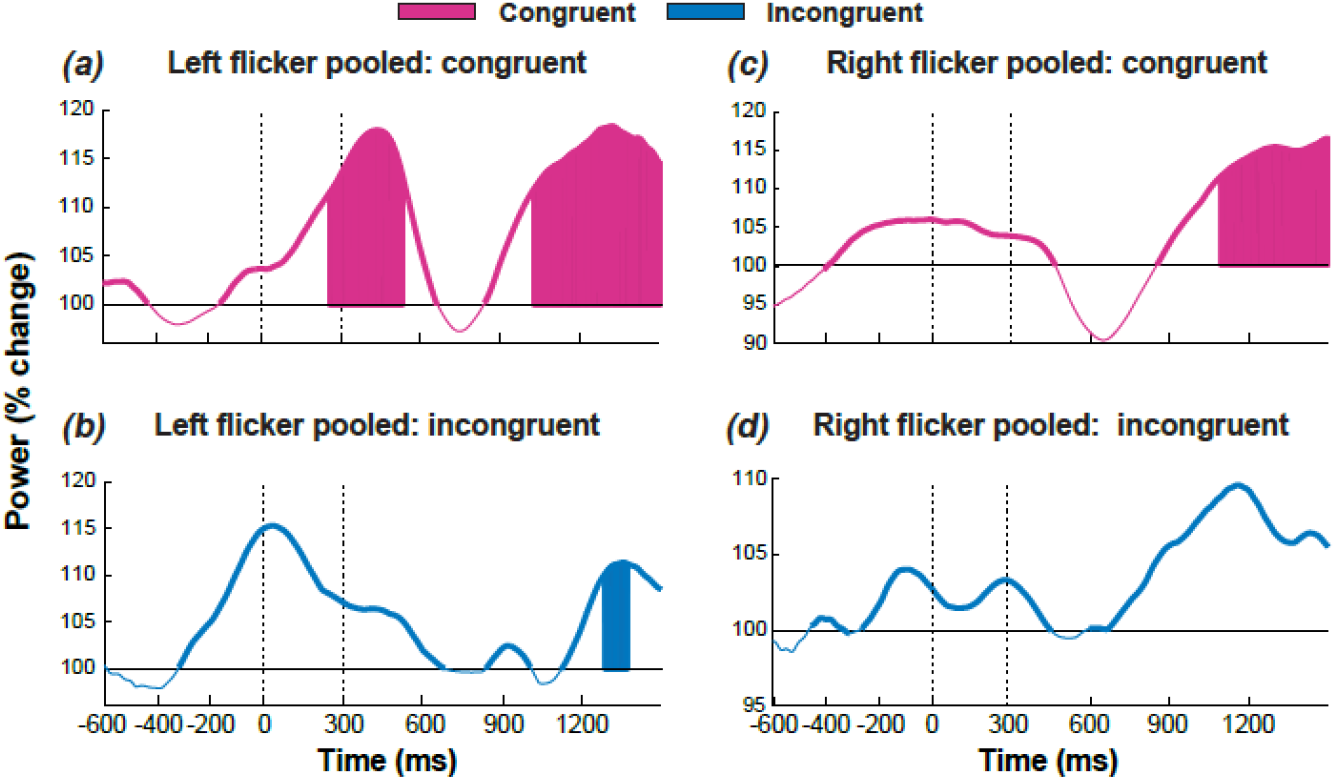
SSVEP changes in amplitude for the left (a, b) and right flicker (c, d) stimuli pooled over number magnitude for (a, c) congruent and (b, d) incongruent conditions. Thicker lines represent when changes are above zero from baseline and thinner line represent changes below baseline. *y* axis corresponds to SSVEP percentage change amplitudes. Shaded areas represent statistically significant temporal differences between the observed increasing in the SSVEP and the baseline (i.e., 100%). Dotted line at time 0 represents time stimulus onset and dotted line at 300 ms represents the stimulus offset. *p* <.05

Thus, initially we pooled all the conditions by congruency and flicker factors to identify any changes in the SSVEP amplitude of the flicker frequencies. For the left and right visual field flicker we observed statistically significant changes above the baseline for the congruent and incongruent conditions. Regarding the left flicker congruent conditions, a significant increasing in SSVEP amplitude was observed during the time of stimulus presentation and reaching its higher peak at 432 ms after stimulus onset and decreasing again until reaching the 100% (i.e., baseline). And from 658 ms to the end of the data epoch the SSVEP amplitude showed a significant increasing again. Additionally, for the left flicker congruent conditions two different time windows showed significant changes with respect to the baseline. The first-time window was between 250 and 530 ms and the second time window was between 1017 ms to the end of the data epoch (Figure 8a). For the left flicker incongruent conditions, a significant increasing in SSVEP amplitude was observed from before the stimulus onset and gradually decreasing until reach the 100% at 682 ms. Then, two more significant changes were obtained one between 837 and 1010 ms and the second one from 1125 ms to the end of the data epoch. For the incongruent conditions the statistically significant time window difference between the SSVEP amplitude with respect to the baseline was observed between 1285 and 1375 ms (Figure 8b).

For the right flicker congruent conditions significant SSVEP amplitude changes with respect to the baseline were observed before stimulus onset from -391 ms to 465 ms, and from 852 ms to the end of the data epoch. The statistically significant time window difference between the SSVEP changes with respect to the baseline was observed between 1092 ms to the end of the data epoch (Figure 8c). For the right flicker incongruent conditions significant changes in the SSVEP amplitude were obtained from -274 ms before stimulus onset to 451 ms and after stimulus offset and from 583 ms to the end of the data epoch. For the right flicker incongruent conditions there were no statistically significant time windows differences between the baseline and the SSVEP changes in amplitude (Figure 8d).

The observed significant changes in the SSVEP amplitude for both flicker frequencies reflect the attention modulation effects resulting from observation of numbers while participants classified them as odd or even. Note however that significant changes in the SSVEP amplitude observed before stimulus onset correspond to the time period used for baseline normalization.

Then, we proceeded to evaluate the dynamic SSVEP for each of the congruent and incongruent conditions, separately for the left and right flicker (all *p* values <.05, cluster-corrected for multiple correlated tests over time points). For the left flicker when small numbers (1,2,3,4) where displayed we did not obtain any statistically significant differences in the SSVEP amplitude between the congruent and incongruent conditions after the stimulus presentation (Figure 9a).

**Figure 9.**
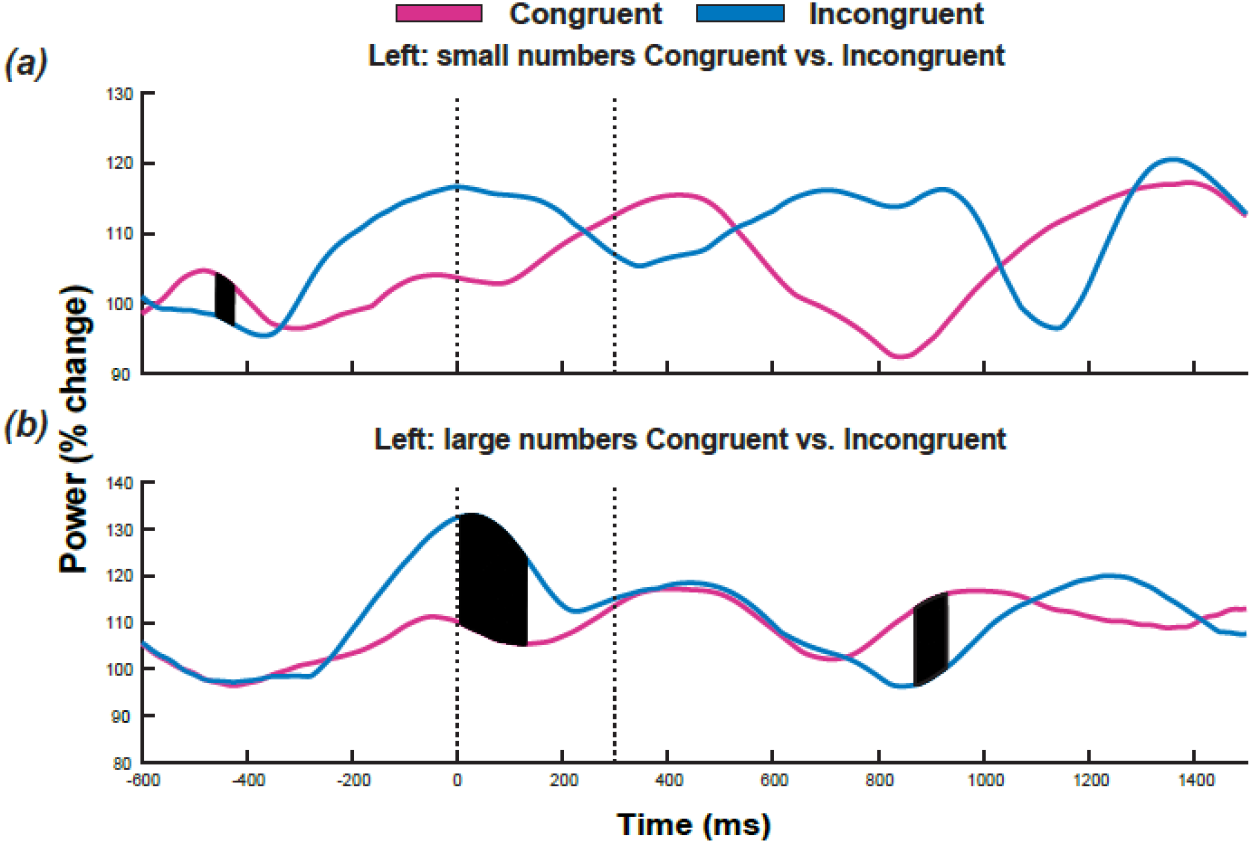
Dynamic SSVEP for the flicker frequency to the left (20 Hz) for small (a) and large magnitude numbers (b). Pre stimulus time between −600 to −200 ms corresponds to the baseline-normalization period. *y* axes correspond to SSVEP percentage change amplitudes. Dotted line at time 0 represents time stimulus onset and dotted line at 300 ms represents the time stimulus offset. Shaded areas represent statistically significant differences between conditions. Notice that the shaded area representing statistically significant differences (from −462 to −417 ms) corresponds to the time window in the baseline period used for normalization. *p* < .05

On the contrary, when large numbers were displayed (6,7,8,9) significant effects were observed at two different time windows (Figure 9b) between the congruent and incongruent conditions. First, increasing SSVEP amplitude for the time window between 0 and 125 ms was observed for the incongruent conditions, but after stimulus presentation the increasing in the SSVEP amplitude was observed for the congruent conditions in the time interval between 868 and 931 ms. Figure 10b shows the results of the nonparametric permutation testing that was used in combination with cluster correction to evaluate condition differences in the dynamic SSVEP over time for correct trials and the left flicker frequency.

**Figure 10.**
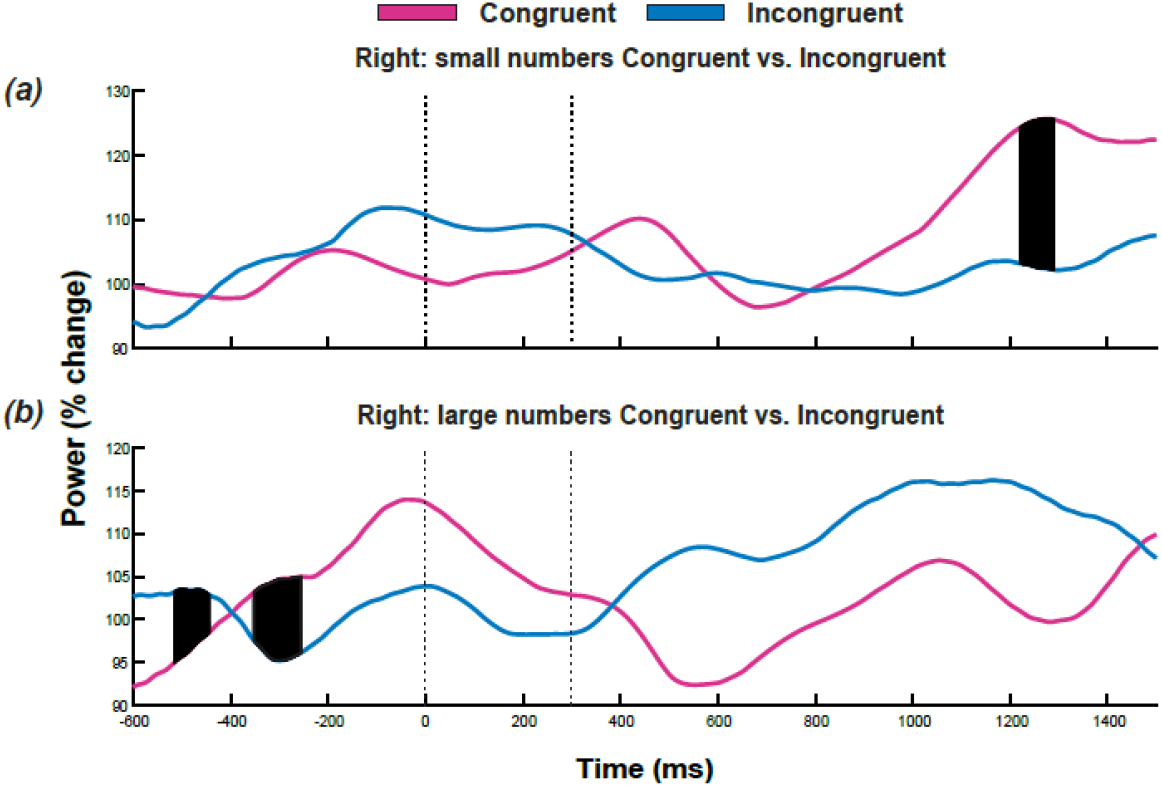
Results for the flicker frequency to the right (24 Hz) for the small (a) and large magnitude numbers (b). Arrangement of Figure 4.9 is the same as in Figure 4.8. Notice that the shaded areas representing statistically significant differences (in the time windows between −513 and −49 and −345 to −254 ms) correspond to the time window in the baseline period used for normalization. *p* < .05

For the flicker stimuli on the right, statistically significant effects in the SSVEP amplitude were observed for congruent trials in the time window between 1225 ms and 1284 ms after stimulus offset as compared with incongruent trials when small numbers stimuli were presented (Figure 10a). Regarding trials displaying large numbers no significant effects in the change of the SSVEP amplitude were observed between the congruent and incongruent conditions (Figure 10b). Figure 10 shows the results of the nonparametric permutation testing that were used in combination with cluster correction to evaluate condition differences in dynamic SSVEP over time for correct trials for the right flicker.

## DISCUSSION

In this study we have used the parity judgment task to induce the so-called SNARC effect and to evaluate whether the simple observation of numbers, when their magnitude is irrelevant, has an attention-modulation effect in the SSVEP amplitude producing the so-called SNARC-spatial attention effect. We implemented a parity judgment task because it is the most frequently task used to study the SNARC effect (Wood, Willmes, Nuerk, & Fischer, 2008), allowing the evaluation of possible implicit spatial attention elicited by seeing the numbers. In our design, participants classified a number as odd or even with the hand response mapping left for odd and right hand for even numbers. We observed significant changes in the SSVEP amplitude with respect to the baseline for the left (20 Hz) and right (24 Hz) flicker for both, the congruent and incongruent conditions. And statistically significant differences between the congruent and incongruent conditions were larger for the congruent conditions for the flicker stimuli on the left. Our results support the interaction between numbers and space as described by the SNARC effect (Dehaene et al., 1993; Fias & Fischer, 2004a) and provide psychophysiological evidence for the hypothesis that observing numbers, even if their magnitude is irrelevant (i.e., parity task), has an effect in the automatic allocation of spatial attention, the so-called SNARC-spatial attention effect.

Our study is the first approach to evaluate the SNARC effect using the SSVEP in a parity judgment task. Behaviorally we have shown that indeed our task induced the SNARC effect improving performance for the congruent versus the incongruent conditions. Our results are in line with previous studies (Chinello et al., 2012; Göbel et al., 2006; Schwarz & Keus, 2004) confirming the positive effect between mental representation of space and numbers while performing a task that involves numbers, even if their magnitude is not relevant for the task (i.e., the parity task). We observed a positive interaction between numbers magnitude and hand response in reaction time and proportion of correct responses, meaning that for small, odd numbers the response time and accuracy performance was better when responded with the left hand as compared with large odd numbers and left hand. And similar performance was observed for large even numbers when responded with the right hand as compared with small even numbers. These results are also consistent with the hypotheses that the way numbers are mentally represented have an effect on performance when there is a relation between the magnitude of the number and the spatial response. And we interpret our results as supporting evidence for the interaction between numbers and space as explained by the MNL and also by the SRC model, but not exclusively one or the other, which is consistent with Santens & Gevers (2008) who claimed that the SNARC effect does not necessarily reflect the mapping between number magnitude and response positions, but that there is an intermediate stage where stimuli are categorized producing a preferential mapping that relies on linguistic features. It is beyond the scope of this report to include comprehensive assessments of the various SNARC effect theories, but readers are directed to (Fias & Fischer, 2004b) and (Bonato et al., 2012) for an extended review of the SNARC effect.

### SSVEP results and the SNARC effect

In this study we evaluated whether numbers have an effect allocating spatial attention during a non-spatial attention task and we included the SSVEP in our paradigm to elucidate if that could be the case in the parity judgment task. Because it has been shown that SSVEP amplitude increases as a function of attention (e.g., Andersen & Müller, 2010; Ding, Sperling, & Srinivasan, 2006; Müller et al., 1998) even if the flicker is irrelevant for the task (Hillyard et al., 1997), we expected changes in the SSVEP amplitude to be larger for congruent as compared with the incongruent conditions. Our results provided evidence supporting the hypothesis that observation of numbers – even without actively processing their magnitude – induced the SNARC-spatial attention effect during the parity judgment task. We observed contralateral SSVEP topographical responses for each of the flickering stimuli at posterior sites (Figure 6) that are consistent with the observed hemifield response in a regular visuospatial attention paradigm (Corbetta & Shulman, 2002; Hopfinger, Buonocore, & Mangun, 2000). And the SSVEP responses for congruent conditions showed significant changes in the amplitude as compared with incongruent conditions after stimulus offset (Figures 8 and 10). These results support the hypothesis that number observation induces allocation of spatial attention, meaning that numbers boost SSVEP amplitude being larger for the hemifield cued by number magnitude. The observed changes in the SSVEP amplitude are consistent with the attention effects previously described in spatial attention (Morgan, Hansen, & Hillyard, 1996; Müller, Malinowski, Gruber, & Hillyard, 2003); and, since in our task participants were requested to ignore the flickering stimulus, the observed effects can be attributed to the allocation of attention to one specific visual hemifield when the observed number is spatially related to the attended hemifield (Fischer et al., 2003) and its position in the MNL theory (Dehaene et al., 1993).

It has been proposed that the SNARC effect is evidence for the automatic activation of number magnitude even when the magnitude is irrelevant for the task (Dehaene et al., 1993; Schwarz & Heinze, 1998) as in the parity task. To further evaluate the effect between large and small numbers in the allocation of spatial attention, we evaluated the difference between congruent and incongruent conditions for the small and large numbers separately for each flicker frequency. For the SSVEP response in the left hemifield, trials displaying large magnitude numbers showed significant increasing of the SSVEP amplitude for congruent trials as compared with incongruent trials and no significant differences were observed for the left flicker when small numbers were display. Similarly, the SSVEP response in the right hemifield showed a significant effect for the congruent conditions when small magnitude numbers were displayed. These results confirm that, although number’s magnitude was not relevant for the task it is possible that numeric semantic information was activated (Sandrini & Rusconi, 2009), which in turn may have induced a shift in spatial allocation of attention. This is consistent with the contralateral hemispheric activation in visual tasks as previously described (Ranzini et al., 2009; Salillas, El Yagoubi, & Semenza, 2008). Our results provide physiological evidence for the hypothesis that observing numbers has an effect on spatial attention, driving covert spatial attention toward the position of number’s magnitude in the MNL while making a decision possibly because during the classification of the number as odd or even its number semantic is activated.

### SNARC effect and hand mapping response

Our study is the first approach, to our knowledge, to evaluate the SNARC effect using the SSVEP, and it could be argued that using only one response mapping we didn’t control for the effects of the responding hand. However, previous studies have shown that the SNARC effect is determined by the spatial position of the response (e.g., number magnitude) and not by the responding hand when using crossed-over hands (left hand pressing right button; Dehaene et al., 1993, Experiment 6), or during unimanual tasks (Fischer, 2003). Similarly, Dehaene et al. (1993) reported the presence of the SNARC effect for left-handed participants (Experiment 5) and during crossed hands experimental conditions on the response button (Experiment 6). Then, our behavioral results and the individual analysis of performance between hand response and number stimulus are consistent with previous studies and support the validity of our experimental design regarding the non-counterbalancing of hand mapping (Gut et al., 2012). Nonetheless, we strongly recommend counterbalancing the hand response mapping in future SNARC-SSVEP studies in order to elucidate the possible effects of different response mappings.

## CONCLUSION

This study has provided behavioral and EEG evidence for the effects of the mental representation of numbers, the mental representation of space, and the interaction between them during a parity task improving performance. On the other hand, we have also provided some relevant evidence for the use of the SSVEP technique to study and better understand the SNARC effect, mental representation of numbers, and how these two processes interact to have an effect on cognition while observing numbers. Our results are in line with different cognitive models explaining the SNARC effect (e.g., MNL, the SRC, dual-route or the PCT models) and support the hypothesis that when numbers are part of the task, but their magnitude is not task-relevant, number semantics are activated having an effect on allocating spatial attention, the so-called SNARC-spatial attention effect. Therefore, the SNARC-spatial attention effect as shown here is present when numbers are relevant for the task. The EEG SSVEP evidence we obtained supports the thesis that the SNARC effect is a cognitive effect resulting from the mental representation of numbers and its relationship with the space representation, more than merely a motor effect. Nonetheless, in order to further confirm our results and/or provide stronger evidence of the SNARC effect in the SSVEP amplitude results during a parity judgment task, some modifications of the paradigm and the task are recommended.

## ACKNOWLEDGEMENTS

A. M. C. is funded by Departamento Administrativo de Ciencia, Tecnología e Innovación, Colciencias call 529/ 2011 (Colombia). M. X. C. is funded by an ERC-StG 638589. R.G. is funded by an ERC awarded to Dr. Rufin Vanrullen (CerCo institute, Tolouse, France).

## AUTHOR CONTRIBUTIONS

M.X.C. and R.G. conception and design of research; M.X.C. analysed data; A.M.C. performed experiments; analysed data; M.X.C. and A.M.C. interpreted results of experiments; A.M.C. prepared figures; drafted manuscript; A.M.C., M.X.C. and R.K.R. edited and revised manuscript; M.X.C. and R.K.R. approved final version of manuscript.

## REFERENCES

Andersen, S. K., & Müller, M. M. (2010). Behavioral performance follows the time course of neural facilitation and suppression during cued shifts of feature-selective attention. Proceedings of the National Academy of Sciences of the United States of America, 107(31), 13878–82. http://doi.org/10.1073/pnas.1002436107

Andres, M., Ostry, D. J., Nicol, F., & Paus, T. (2008). Time course of number magnitude interference during grasping. Cortex, 44(4), 414–419. http://doi.org/10.1016/j.cortex.2007.08.007

Bonato, M., Zorzi, M., & Umiltà, C. (2012). When time is space: Evidence for a mental timeline. Neuroscience and Biobehavioral Reviews, 36(10), 2257–2273. http://doi.org/10.1016/j.neubiorev.2012.08.007

Brainard, D. H. (1997). The Psychophysics Toolbox. Spatial Vision, 10(4), 433–436. http://doi.org/10.1163/156856897X00357

Chinello, A., de Hevia, M. D., Geraci, C., & Girelli, L. (2012). Finding the spatial-numerical association of response codes (SNARC) in signed numbers: Notational effects in accessing number representation. Functional Neurology, 27(3), 177–185.

Cohen, M. X. (2014). Analyzing Neural Time Series: Theory and practice (First). Cambridge, MA: MIT Press. http://doi.org/10.1017/CBO9781107415324.004

Cohen, M. X., & Gulbinaite, R. (2017). Rhythmic entrainment source separation: Optimizing analyses of neural responses to rhythmic sensory stimulation. NeuroImage, 147(August 2016), 43–56. http://doi.org/https://dx.doi.org/10.1016/j.neuroimage.2016.11.036

Corbetta, M., & Shulman, G. L. (2002). Control of goal-directed and stimulus-driven attention in the brain. Nature Reviews Neuroscience, 3(3), 201–215. http://doi.org/10.1038/nrn755

Dehaene, S., Bossini, S., & Giraux, P. (1993). The Mental representation of Parity and Number Magnitude. Journal of Experimental Psychology. General, 122(3), 317–396

Delorme, A., & Makeig, S. (2004). EEGLAB: an open-source toolbox for analysis of single-trial EEG dynamics including independent component analysis. Journal of Neuroscience Methods, 134(1), 9–21. http://doi.org/10.1016/j.jneumeth.2003.10.009

Ding, J., Sperling, G., & Srinivasan, R. (2006). Attentional modulation of SSVEP power depends on the network tagged by the flicker frequency. Cerebral Cortex, 16(7), 1016–29. http://doi.org/10.1093/cercor/bhj044

Doherty, J. R., Rao, A., Mesulam, M. M., & Nobre, A. C. (2005). Synergistic effect of combined temporal and spatial expectations on visual attention. The Journal of Neuroscience, 25(36), 8259–8266. http://doi.org/10.1523/JNEUROSCI.1821-05.2005

Fias, W., Brysbaert, M., Geypens, F., & d’Ydewalle, G. (1996). The importance of magnitude information in numerical processing: Evidence from the SNARC effect. Mathematical Cognition, 2(1), 995–110.

Fias, W., & Fischer, M. (2004a). Spatial representation of number. Handbook of Mathematical Cognition. http://doi.org/10.4324/9780203998045.ch3

Fias, W., & Fischer, M. H. (2004b). Spatial Representation of numbers. In J. I. D. Campbell (Ed.), Handbook of Mathematical Cognition (pp. 43–54). Routledge. http://doi.org/10.4324/9780203998045.ch3

Fias, W., Lauwereyns, J., & Lammertyn, J. (2001). Irrelevant digits affect feature-based attention depending on the overlap of neural circuits. Cognitive Brain Research, 12, 415–423.

Fischer, M. H. (2003). Spatial representations in number processing - Evidence from a pointing task. Visual Cognition, 10(4), 493–508. http://doi.org/10.1080/13506280244000186

Fischer, M. H., Warlop, N., Hill, R. L., & Fias, W. (2004). Oculomotor Bias Induced by Number Perception. Experimental Psychology, 51(2), 91–97. http://doi.org/10.1027/1618-3169.51.2.91

Fitts, P. M., & Seeger, C. M. (1953). S-R compatibility: Spatial characteristics of stimulus and response codes. Journal of Experimental Psychology, 46(3), 199–210.

Galfano, G., Rusconi, E., & Umiltà, C. (2006). Number magnitude orients attention, but not against one’s will. Psychonomic Bulletin & Review, 13(5), 869–74. http://doi.org/10.3758/BF03194011

Gevers, W., Caessens, B., & Fias, W. (2005). Towards a common processing architecture underlying Simon and SNARC effects. European Journal of Cognitive Psychology, 17(5), 659–673. http://doi.org/10.1080/09541440540000112

Gevers, W., Ratinckx, E., De Baene, W., & Fias, W. (2006). Further evidence that the SNARC effect is processed along a dual-route architectures: Evidence from the lateralized readiness potential. Experimental Psychology, 53(1), 58–68. http://doi.org/10.1027/1618-3169.53.1.58

Göbel, S. M., Calabria, M., Farnè, A., & Rossetti, Y. (2006). Parietal rTMS distorts the mental number line: Simulating “spatial” neglect in healthy subjects. Neuropsychologia, 44(6), 860–868. http://doi.org/10.1016/j.neuropsychologia.2005.09.007

Gut, M., Szumska, I., Wasilewska, M., & Jaśkowski, P. (2012). Are low and high number magnitudes processed differently while resolving the conflict evoked by the SNARC effect? International Journal of Psychophysiology, 85(1), 7–16. http://doi.org/10.1016/j.ijpsycho.2012.02.007\

Herrmann, C. S. (2001). Human EEG responses to 1-100 Hz flicker: Resonance phenomena in visual cortex and their potential correlation to cognitive phenomena. Experimental Brain Research, 137(3–4), 346–353. http://doi.org/10.1007/s002210100682

Hesse, P. N., & Bremmer, F. (2017). The SNARC effect in two dimensions: Evidence for a frontoparallel mental number plane. Vision Research, 130, 85–96. http://doi.org/10.1016/j.visres.2016.10.007

Hillyard, S. A., Hinrichs, H., Tempelmann, C., Morgan, S. T., Hansen, J. C., Scheich, H., & Heinze, H. J. (1997). Combining steady-state visual evoked potentials and f MRI to localize brain activity during selective attention. Human Brain Mapping, 5(4), 287–292. http://doi.org/10.1002/(SICI)1097-0193(1997)5:4<287::AID-HBM14>3.0.CO;2-B

Hopfinger, J. B., Buonocore, M. H., & Mangun, G. R. (2000). The neural mechanisms of top-down attentional control. Nature Neuroscience, 3(3), 284–291. http://doi.org/10.1038/72999

Keus, I. M., Jenks, K. M., & Schwarz, W. (2005). Psychophysiological evidence that the SNARC effect has its functional locus in a response selection stage. Cognitive Brain Research, 24(1), 48–56. http://doi.org/10.1016/j.cogbrainres.2004.12.005

Keus, I. M., & Schwarz, W. (2005). Searching for the functional locus of the SNARC effect: evidence for a response-related origin. Memory & Cognition, 33(4), 681–695. http://doi.org/10.3758/BF03195335

Kleiner, M., Brainard, D. H., & Pelli, D. (2007). “Whats new in Psychtoolbox-3?” In Thirtieth European Conference on Visual Perception (Vol. 36, pp. 1–235). Arezzo.

Kornblum, S., Hasbroucq, T., & Osman, A. (1990). Dimensional overlap: cognitive basis for stimulus-response compatibility-A model and taxonomy. Psychological Review, 97(2), 253–270. http://doi.org/10.1037/0033-295X.97.2.253

Luck, S. J. (2005a). An Introduction to event-related potentials and their neural origins. An introduction to the event-related potential technique. Michigan, MA, USA: MIT Press. http://doi.org/10.1007/s10409-008-0217-3

Maris, E., & Oostenveld, R. (2007). Nonparametric statistical testing of EEG-and MEG-data. Journal of Neuroscience Methods, 164(1), 177–190. http://doi.org/10.1016/j.jneumeth.2007.03.024

Morgan, S. T., Hansen, J. C., & Hillyard, S. A. (1996). Selective attention to stimulus location modulates the steady-state visual evoked potential. Proceedings of the National Academy of Sciences of the United States of America, 93(10), 4770–4. http://doi.org/10.1073/pnas.93.10.4770

Moyer, R. S., & Landauer, T. K. (1967). Time required for Judgements of Numerical Inequality. Nature, 215(5109), 1519–1520. http://doi.org/10.1038/2151519a0

Müller, M. M., Malinowski, P., Gruber, T., & Hillyard, S. A. (2003). Sustained division of the attentional spotlight. Nature, 24, 309–312. http://doi.org/10.1038/nature01744.1.

Müller, D., & Schwarz, W. (2007). Exploring the mental number line: Evidence from a dual-task paradigm. Psychological Research, 71(5), 598–613. http://doi.org/10.1007/s00426-006-0070-6

Müller, M. M., Picton, T. W., Valdes-Sosa, P., Riera, J., Teder-Sälejärvi, W. A., & Hillyard, S. A. (1998). Effects of spatial selective attention on the steady-state visual evoked potential in the 20-28 Hz range. Cognitive Brain Research, 6(4), 249–61. http://doi.org/10.1016/S0926-6410(97)00036-0

Pelli, D. G. (1997). The Videotoolbox software for visual psychophysics: Transforming numbers into movies. Spatial Vision. http://doi.org/10.1163/156856897X00366

Pinel, P., Piazza, M., Le Bihan, D., & Dehaene, S. (2004). Distributed and overlapping cerebral representations of number, size, and luminance during comparative judgments. Neuron, 41(6), 983–993. http://doi.org/10.1016/S0896-6273(04)00107-2

Posner, M. I. (1980). Orienting of attention. Quarterly Journal of Experimental Psychology, 32(1), 3–25. http://doi.org/10.1080/00335558008248231

Posner, M. I., Snyder, C. R., & Davidson, B. J. (1980). Attention and the detection of signals. Journal of Experimental Psychology: General, 109(2), 160–174. http://doi.org/10.1037/0096-3445.109.2.160

Proctor, R. W., & Cho, Y. S. (2006). Polarity correspondence: A general principle for performance of speeded binary classification tasks. Psychological Bulletin, 132(3), 416–442. http://doi.org/10.1037/0033-2909.132.3.416

Ranzini, M., Dehaene, S., Piazza, M., & Hubbard, E. M. (2009). Neural mechanisms of attentional shifts due to irrelevant spatial and numerical cues. Neuropsychologia, 47(12), 2615–24. http://doi.org/10.1016/j.neuropsychologia.2009.05.011

Regan, D. (1989). Human brain electrophysiology: Evoked potentials and evoked magnetic fields in science and medicine (Vol. 1). New York: Elsevier.

Restle, F. (1970). Speed of adding and comparing numbers. Journal of Experimental Psychology, 83(2, Pt.1), 274–278. http://doi.org/10.1037/h0028573

Ristic, J., Wright, A., & Kingstone, A. (2006). The number line effect reflects top-down control. Psychonomic Bulletin & Review, 13(5), 862–868. http://doi.org/10.3758/BF03194010

Salillas, E., El Yagoubi, R., & Semenza, C. (2008). Sensory and cognitive processes of shifts of spatial attention induced by numbers: an ERP study. Cortex; a Journal Devoted to the Study of the Nervous System and Behavior, 44(4), 406–13. http://doi.org/10.1016/j.cortex.2007.08.006

Sandrini, M., & Rusconi, E. (2009). A brain for numbers. Cortex, 45(7), 796–803. http://doi.org/10.1016/j.cortex.2008.09.002

Santens, S., & Gevers, W. (2008). The SNARC effect does not imply a mental number line. Cognition, 108(1), 263–270. http://doi.org/10.1016/j.cognition.2008.01.002

Schwarz, W., & Heinze, H. J. (1998). On the interaction of numerical and size information in digit comparison: A behavioral and event-related potential study. Neuropsychologia, 36(11), 1167–1179. http://doi.org/10.1016/S0028-3932(98)00001-3

Schwarz, W., & Keus, I. M. (2004). Moving the eyes along the mental number line: Comparing SNARC effects with saccadic and manual responses. Perception & Psychophysics, 66(4), 651–664. http://doi.org/10.3758/BF03194909

Schwarz, W., & Müller, D. (2006). Spatial associations in number-related tasks: A comparison of manual and pedal responses. Experimental Psychology, 53(1), 4–15. http://doi.org/10.1027/1618-3169.53.1.4

Shaki, S., Fischer, M. H., & Göbel, S. M. (2012). Direction counts: A comparative study of spatially directional counting biases in cultures with different reading directions. Journal of Experimental Child Psychology, 112(2), 275–281. http://doi.org/10.1016/j.jecp.2011.12.005

van Dijck, J. P., Abrahamse, E. L., Acar, F., Ketels, B., & Fias, W. (2014). A working memoryaccount of the interaction between numbers and spatial attention. Quarterly Journal of Experimental Psychology, 67(8), 1500–1513. http://doi.org/10.1080/17470218.2014.903984

Vogel, S. E., Grabner, R. H., Schneider, M., Siegler, R. S., & Ansari, D. (2013). Overlapping and distinct brain regions involved in estimating the spatial position of numerical and non-numerical magnitudes: An fMRI study. Neuropsychologia, 51(5), 979–989. http://doi.org/10.1016/j.neuropsychologia.2013.02.001

Weis, T., Estner, B., van Leeuwen, C., & Lachmann, T. (2016). SNARC (spatial–numerical association of response codes) meets SPARC (spatial–pitch association of response codes): Automaticity and interdependency in compatibility effects. The Quarterly Journal of Experimental Psychology, 69(7), 1366–1383. http://doi.org/10.1080/17470218.2015.1082142

Wood, G., Willmes, K., Nuerk, H.-C., & Fischer, M. H. (2008). On the cognitive link between space and number: a meta-analysis of the SNARC effect. Psychology Science Quarterly, 4(4), 489–525. http://doi.org/10.1027/1618-3169.52.3.187

Zanolie, K., & Pecher, D. (2014). Number-induced shifts in spatial attention: A replication study. Frontiers in Psychology, 5(AUG),1–10. http://doi.org/10.3389/fpsyg.2014.00987

Zohar-shai, B., Tzelgov, J., Karni, A., Rubinsten, O., Zohar-shai, B., & Tzelgov, J. (2017). It DoesExist ! A Left-to-Right Spatial –Numerical Association of Response Codes (SNARC) Effect Among Native Hebrew Speakers. Journal of Experimental Psychology: Human Perception and Performance, 43(4), 719–728. http://doi.org/10.1037/xhp0000336

Zorzi, M., Priftis, K., & Umiltà, C. (2002). Brain damage: Neglect disrupts the mental number line. Nature, 417(6885), 138–139. http://doi.org/10.1038/417138a

